# Respiratory syncytial virus matrix protein assembles as a lattice with local and extended order that coordinates the position of the fusion glycoprotein

**DOI:** 10.1101/2021.10.13.464285

**Authors:** Bryan S. Sibert, Joseph Y. Kim, Jie E. Yang, Zunlong Ke, Christopher C. Stobart, Martin M. Moore, Elizabeth R. Wright

## Abstract

Respiratory syncytial virus (RSV) is a significant cause of respiratory illness in young children and adults worldwide. There is currently no vaccine or targeted antiviral for RSV. RSV is an enveloped, filamentous, negative-strand RNA virus. Individual virions vary in both diameter and length, with an average diameter of ∼130 nm and ranging from ∼500 nm to over 10 μm in length. The RSV matrix (M) protein is peripherally associated with the interior of the viral membrane. Though the general arrangement of structural proteins within the virion is known, the molecular organization of M and other structural proteins was previously unknown. Here, using whole-cell cryo-electron tomography and sub-tomogram averaging, we show that M is arranged in a packed helical-like lattice of M-dimers ordered at an angle of ∼47° to the viral long axis. Sub-tomogram averages including F and M indicate that the position of F on the viral surface is correlated with the underlying M lattice. Finally, we report that RSV F is frequently observed as pairs, with the F trimers oriented in an anti-parallel conformation to support potential interaction between trimers. These results provide insight into RSV assembly and virion organization and may aid in the identification and development of RSV vaccines and anti-viral targets.

## Introduction

Respiratory syncytial virus (RSV) is a negative-strand RNA virus in the order *Mononegavirales*, which includes many pathogenic viruses such as measles virus, rabies virus, and Ebola virus that can cause severe disease and death [1]. RSV causes respiratory illness in both children and adults and is estimated to cause over three million hospitalizations annually [2, 3]. Though there are several ongoing vaccine clinical trials [4], there is currently no approved vaccine or virus-specific treatment for RSV. Prophylactic treatment with a monoclonal antibody targeting the RSV F protein is effective to prevent severe illness in high-risk children [5].

RSV virions are enveloped, filamentous, and highly pleiomorphic in nature. Virions can range in length from less than one to over ten micrometers and virions also vary in diameter, with an average diameter of 130 nm [6]. Virion morphology and fitness may be altered by heat or mechanical stress during virus purification [7, 8]. RSV has three surface proteins, fusion glycoprotein (F), attachment glycoprotein (G), and small hydrophobic protein (SH). RSV F is a class I fusion protein present as a homotrimer on the viral surface. The F protein is a promising target for vaccine development due to its sequence conservation between strains and the prevalence of neutralizing antibodies against prefusion-F [9-11]. Structures for purified pre- and post-fusion F-trimers have been solved using x-ray crystallography and cryo-EM single particle analysis [10, 12, 13]. A number of structures and conformational states of RSV-F on intact virions have been determined, to a lower resolution, using cryo-ET and sub-tomogram averaging (STA) [8, 14]. RSV-G is the primary attachment protein for RSV, although it has been shown to be non-essential for infection in some cell lines [15]. G can mediate attachment through heparin sulfate and the protein CX3CR1 [16]. RSV SH is a small hydrophobic protein that has been shown to have viroporin activity [17]. SH is not required for viral entry or replication [18].

RSV matrix (M) is a peripheral membrane-protein lining the majority of the interior of the inner viral membrane [6, 14, 19]. The presence of matrix, or a functionally similar protein, is widely conserved amongst negative-strand RNA viruses. Many studies of these viruses have identified a central role for matrix or matrix-like proteins in virion organization and assembly [20-23]. Consistent with this, RSV M is essential for assembly of filamentous virions and virus-like particles (VLPs) [24]. Purified RSV M has been crystallized as both a monomer and a dimer leaving some uncertainty regarding M organization in virions [25, 26]. However, assembly of filamentous virions or virus-like particles (VLPs) is inhibited by mutations that disrupt M dimerization [25]. We previously published results from cryo-ET studies and sub-tomogram averaging of RSV virions indicating that M is arranged in a lattice of M dimers [27, 28]. Assembly of a membrane-associated matrix or matrix-like protein lattice has been shown for several other viruses [29-35]. While RSV M alone is not sufficient for filamentous VLP assembly, only RSV M, F, and P (phosphoprotein) are required [36].

RSV P is a non-catalytic phosphoprotein that is essential for viral RNA synthesis. It’s exact role in virion or VLP assembly remains unknown, but it is known to interact directly with M [37]. P interacts with other structural proteins as well, including L and M2-1, and may be involved in mediating interaction between M and other proteins [36-38]. RSV M associates with the nucleoprotein (N) through mutual neighboring interactions with M2-1 [39]. Cryo-ET and super-resolution fluorescence microscopy demonstrated that M2-1 was present as regularly spaced densities between the M layer and the RNP in virions [7, 14]. In addition to its structural role, M2-1 acts as an ant-termination factor during RNA transcription [40]. RSV N is associated with the genomic RNA in a left-handed helical nucleocapsid [41] that can be observed throughout the virion interior. The RNA-dependent RNA polymerase, L, is associated with N in the viral filaments [42].

In order to study the native structure of RSV and other enveloped viruses, we and others use whole-cell cryo-ET [19, 29, 43-45]. Cells are grown and infected with RSV directly on TEM grids [6, 46, 47]. The grids are then plunge-frozen to rapidly cryo-preserve the cells, associated viruses, and released virions in vitreous ice. By whole-cell cryo-ET, RSV virions are predominantly filamentous; however, irregular, branched, and bent viruses may still be observed [6, 7]. Using cryo-ET and sub-tomogram averaging we show that RSV M is arranged in a helical-like lattice along the interior of the viral envelope. The lattice is composed of M-dimers as indicated by the lattice spacing and modeling of an M-dimer crystal structure into the sub-tomogram average. We further show that the position of F on the viral surface is ordered relative to the underlying M lattice. F is frequently observed in pairs on the viral surface and sub-tomogram averaging of F pairs indicates an anti-parallel arrangement of the F trimers with potential quaternary interactions between the two trimers of a bundle. A more complete understanding of RSV M and F interactions and organization is critical to inform future vaccine design and development.

## Results

### RSV F is arranged in rows and pairs on virions

To determine the organization of RSV structural proteins within intact virions, we performed whole-cell cryo-ET of RSV virions released from infected cells that were grown directly on TEM grids. We used conditions similar to those previously outlined by Ke, et al. [6]. Briefly, BEAS-2B cells were cultured directly on gold TEM grids and infected with RSV-A2mK+ at a multiplicity of infection (MOI) of 10. The cells and viruses on the grids were cryo-fixed by plunge-freezing 24 hours post infection. A slice from a representative tomographic volume is shown in Figure 1A. Consistent with previous reports [6, 46, 48], we were able to identify densities attributed to the viral membrane, the fusion (F) glycoprotein, matrix (M) protein, M2-1, and ribonucleoprotein (RNP) complex (Figure 1B). RSV F was present as densities decorating the surface of the outer viral membrane. The M protein was closely associated with the membrane and appears as a continuous layer beneath the membrane in views such as Figure 1B. Interior to that was a layer of regularly spaced densities that has been attributed to M2-1 [7], though the presence of additional macromolecules in this layer cannot be ruled out. The RNP was frequently observed running along the M2-1 layer as seen in Figure 1B and in the density profile plot in Figure 1C. However, the RNP was not always closely associated with the M2-1 layer and was also be found throughout the interior of the virus, which was consistent with our previously published results [7].

**Figure 1.**
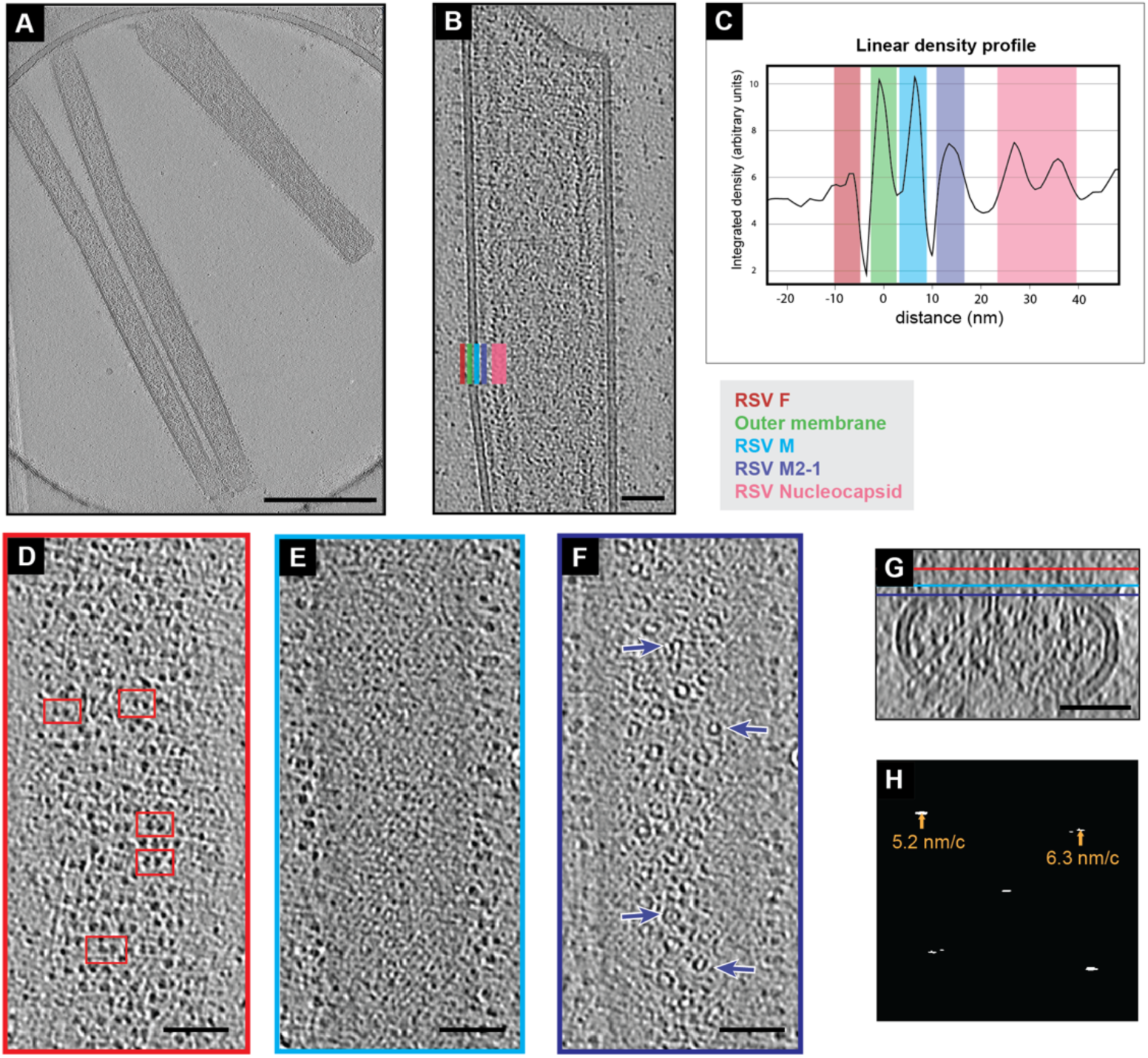
Organization of RSV structural proteins. A) Single z-slice from a representative cryo-tomogram of RSV virions. Protein density is black. B) A reoriented Z-projection (9 nm thick) from a tomogram of an RSV virion. In this orientation, the viral membrane and several structural proteins are localized within columns in the image as indicated by the colored bars in the bottom left. A color-coded legend is indicated to the lower right. C) An average linear density profile across the left-side of the virion in (B) showing the peaks corresponding with each column of proteins or membrane. Colored boxes correspond to the legend and positions indicated in (B). D) Single slice from the virion in (B) showing RSV F on the top of the virion in the X, Y plane (red bar in (G) indicates position of slice in Z). Select examples of pairs of density are highlighted by red boxes. (E) Single slice from the M protein layer from the virion in (B) (cyan bar in (G)). (F) Single slice from the virion in (B) showing the M2-1 layer (purple bar in (H)). Arrows point to ring like densities. (G) Y-projection (9 nm thick) of X, Z slices of the virion in (B), Z-position of slices in (D), (E), and (F) are indicated by the red, cyan, and purple bars respectively. (H) Amplitude spectrum from of slice in (E) (transform is from cropped segment including only the M lattice). Peaks in the spectrum and the corresponding radius in nanometers/cycle are indicated by arrows. 500 nm scale bar for (A), 50 nm scale bar for (B), (D), (E), (F), and (G).

Upon closer examination of the distribution of RSV F along the surface of the virions, it was clear that F was not present as a tightly packed lattice, however, the distribution did not appear random. Figure 1D shows a tomographic slice including F densities on the viral surface in which F appeared to be roughly organized into rows perpendicular to the long axis of the virus. Within these rows F often occurred as pairs in which two trimers were closer to one-another than neighboring trimers. Examples of the F pairs are highlighted in red boxes in Figure 1D.

### Lattice density is present in M-layer of RSV virions

Within tomographic slices that contained the thin M-layer, as shown in Figure 1E, we observed large regions of lattice-like density extending along the virions. The presence of this layer was more apparent when the virion was reoriented to be flat to the viewing plane allowing a larger surface of the M layer to be viewed within a single section. To confirm our initial assessment of the lattice-like density in this layer we analyzed the amplitude spectra of a cropped region of Figure 1E that included only the M lattice (Figure 1F). We observed clear peaks in the spectrum consistent with the presence of a lattice with spacings of 5.2 nm and 6.3 nm as measured from the center of the peaks. The peaks appeared as short horizontal lines covering frequencies from 5.1 – 5.4 nm/c and 6.2 - 6.9 nm/c, respectively. We then determined the power spectra from the M lattice in eleven more similarly cropped tomographic slices of virion segments and consistently observed peaks corresponding to real space distances of approximately 5.3 ± 0.1 nm and 6.4 ± 0.2 nm (mean ± standard deviation). When first orienting the long axis of the virus vertically, as in Figure 1E, the angle of the peaks was similar among all virions with the 5.3 nm peak at 136.0° ± 2.7° from the x-axis and the 6.4 nm peak at 38.2° ± 3.8° from the x-axis. Together, this indicated the presence of a locally ordered helical-like lattice with a conserved orientation in the M layer.

A regular spacing of ∼12.6 nm has been previously reported for M2-1 [7] and such periodic densities were observed in the layer in Figure 1B (purple bar). Looking at the same distance from the from the membrane, but from the top as in Figure 1G, a number of ring-like structures were observed (purple arrows). There was no obvious organization to the rings. Though some of the rings appeared to be consistent in size and shape with nucleocapsid [41], they were not associated with sections of a longer strand of nucleocapsid. Additionally, variation in the ring diameter was noted, though this could be due, in part, to varied orientation of the rings relative to the tomographic slice.

### Sub-tomogram averaging of M-lattice

We applied sub-tomogram averaging to analyze the organization of RSV M in the lattice. Figure 2A shows a tomographic slice of a virion segment overlaid with model points marking the centers of aligned sub-tomograms included in the average. The lattice organization was seen in the distribution of the points, although small gaps were present throughout. The gaps in model points did not definitively indicate gaps in the M lattice because some points may have been missed during particle selection. Density from the M lattice was clearly seen as a series of peaks and troughs beneath the inner and outer leaflets of the viral membrane in the sub-tomogram average (Figure 2B). The isosurface of the sub-tomogram average (Figure 2C) has been colored by z-height to highlight the various structures with viral membrane in green, the M lattice in cyan, and the M2-1 layer in purple. To validate that the lattice structure observed in the sub-tomogram average was consistent with a lattice of M-dimers, molecular surface models of the M dimer crystal structure (PDB 4v23 [25]) were fit into the average (Figure 2D). The M monomers within each dimer have been bicolored cyan and dark blue to better visualize the organization and packing.

**Figure 2.**
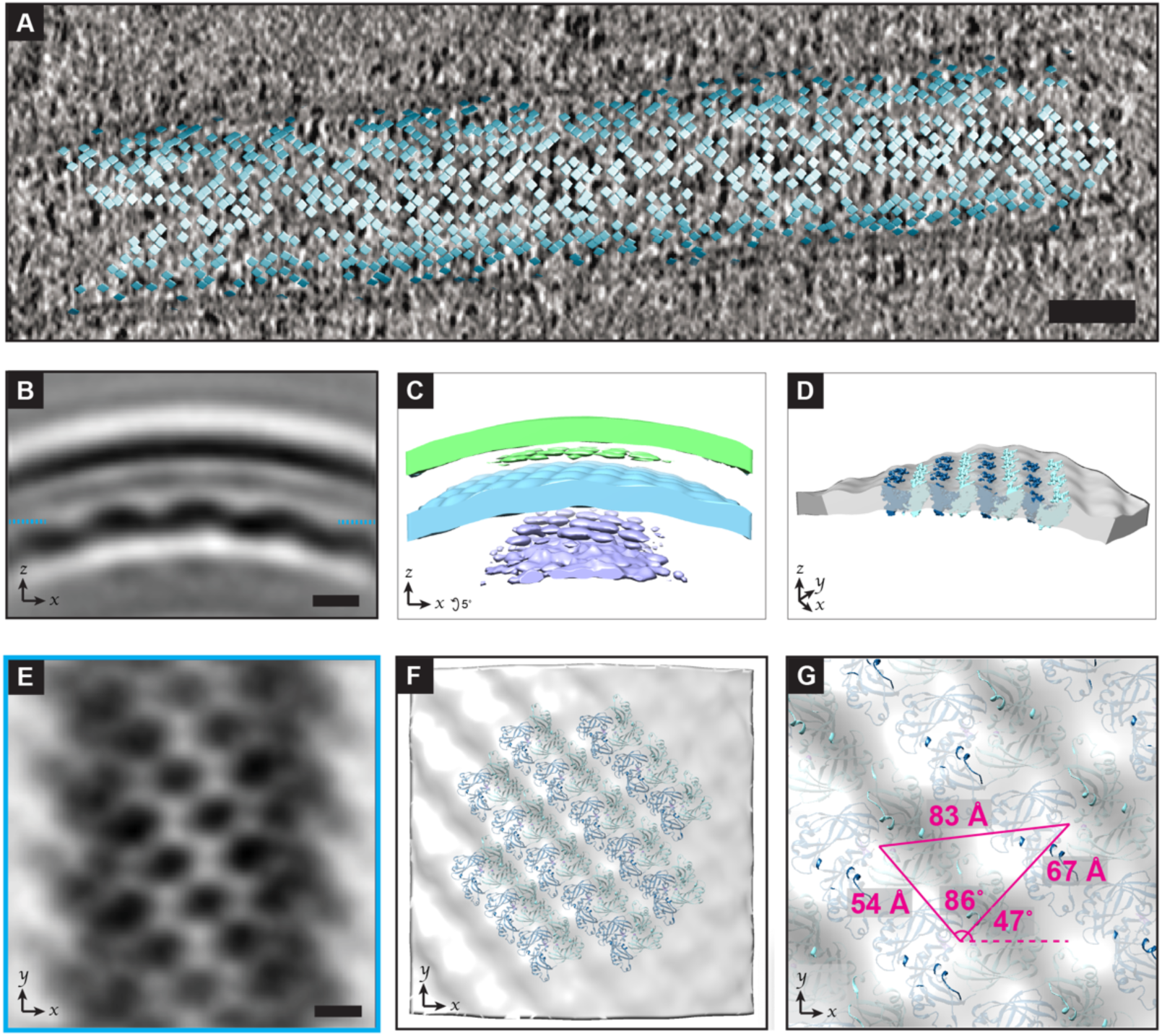
Sub-tomogram average of RSV M. A) Model points for the center of sub-tomograms used in sub-tomogram averaging of RSV M are shown as squares and positioned above a projection of tomographic slices (5.5 nm thick) from the corresponding virion segment. B) Single slice from a sub-tomogram average of RSV M oriented with the M lattice shown underneath the viral membrane. Dashed lines indicate the z-position of the slice in (E). C) Isosurface rendering of RSV M sub-tomogram average including densities from the membrane and underlying M2-1 layer; map is colored by z-height (green - viral membrane, cyan - M layer, purple - M2-1 layer). D) Isosurface of RSV M with model fitting of the M dimer (PDB: 4v23). Models are shown as molecular surfaces with the individual monomers differentially colored (dark blue and cyan). The volume has been rotated and clipped, with non-M densities removed to highlight the structure and model fitting within the M layer. E) Single slice from sub-tomogram average from (B). F) Isosurface of M layer with model fitting of the M-dimer with non-M densities removed, viewed from the membrane side. G) Enlarged view of F overlaid with the average center-to-center distances between individually fit dimers as well as the angles between dimers and the angle relative to the x-axis. (F). 50 nm scale bar for (A), 5 nm scale bar for (B) and (E); volume was lowpass filtered to 21 Å.

The organization and spacing of the lattice was particularly evident when oriented flat in the viewing plane as for the section in Figure 2E. To measure the lattice spacing we used the M-dimer models that were individually fit into the sub-tomogram average using UCSF Chimera (Figure 2D&F). The center to center spacing between the modelled dimers was 5.4 ± 0.7 nm (mean ± standard deviation) and 6.7 ± 1.0 nm. As expected, these values matched closely to those obtained in our analysis of the power spectrum from tomographic slices of M (5.3 ± 0.1 nm and 6.4 ± 0.2 nm). We measured the angles between dimers (as indicated in Figure 2G) to be 85.94 ± 1.29 degrees. The equivalent angle determined from the center of peaks in the power spectrum was 82.3 ± 1.6 degrees. The variations in distances and angles measured within the lattice may be due to factors such as imperfect model fitting or variations in lattice curvature within and between different segments. The range of diameters from the virion segments included in our sub-tomogram average of M was 99 nm – 203.6 nm, with a mean and standard deviation of 135.6 nm and 19.5 nm. (supplemental Figure S4). The potential causes and implications of variation in the lattice spacing are detailed in the discussion section.

### Sub-tomogram averaging of a pair of F trimers

To better understand the organization of F on the virion surface we determined a sub-tomogram average of an F trimer pair. The resulting average contained two trimeric structures that extended approximately 12 nm above the membrane surface, consistent with previously reported structures of prefusion F on virions (Figure 3A&B) [6, 8, 14]. To further validate that the structures were RSV prefusion F homotrimers, we fit in a model from a previously determined crystal structure of RSV F prefusion trimers (PDB: 5w23 [12]) (Figure 3D&E). The sub-tomogram average revealed that the F trimers were not positioned randomly with respect to each other or the virion, but were antiparallel to one another (Figure 3B&E). Furthermore, the placement of an individual trimer was determined by its position within the trimer pair, where one of the vertices of the left most trimer was always pointed in an upward orientation. This positioning was validated by the model fitting as seen in Figure 3D&E. Although we cannot rule out that there may be other arrangements of the trimer clusters, we were unable to generate sub-tomogram averages in which the trimer pairs had other orientation parameters.

**Figure 3.**
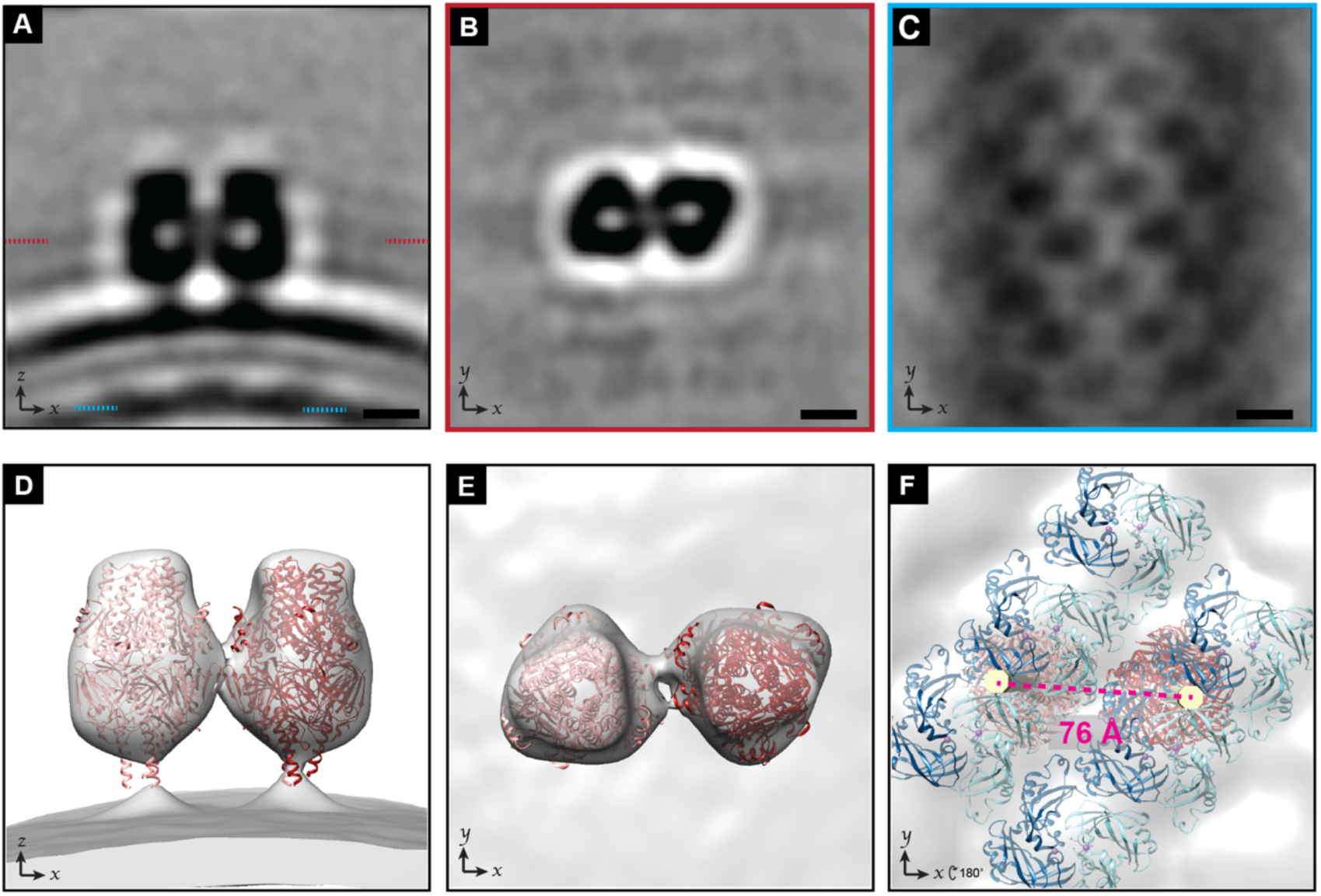
Sub-tomogram average of RSV F pair. A) Slice from a sub-tomogram average of a pair of RSV F trimers on the viral envelope. The outer lipid layer, inner lipid layer, and M layer can be seen underneath. The z-position of the sections in (B) and (C) are indicated with red and cyan dashed lines respectively. B) Slice from the sub-tomogram average from (A), viewed looking towards the membrane from outside of the virus. The two trimers are arranged in an antiparallel fashion. C) Slice from the sub-tomogram average from (A) showing the M layer. D) Isosurface renderings of the sub-tomogram average in (A) with RSV F pre-fusion timer models (PDB: 5w23) fit in. E) Rotated view of (D), viewed looking towards the membrane from outside of the virus. F) Isosurface renderings of the sub-tomogram average in (A) with models of RSV F (PDB: 5w23, red) and models of M dimers (PDB: 4v23, dark blue and cyan) fit in. Viewed from the interior of the virus. A straight rod (yellow) has been modelled in through the center of each F trimer extending down to M. The center-to-center distance between the two fit F-trimers is shown in red. 5 nm scale bar for (A), (B), and (C); volume was lowpass filtered to 24 Å.

### M lattice coordination of F positioning

To better understand the relationship between M and the F glycoproteins on the virion surface, we examined the density corresponding to the M layer within the sub-tomogram average of the F trimer pairs. The lattice organization of the M layer was clearly preserved (Figure 3A&C). If the F pairs were randomly positioned with respect to the underlying M lattice, an average of the M lattice would not have been revealed. Instead, the pair of F trimers and M lattice were resolved within a single sub-tomogram average, which indicated that positioning of M and F are coordinated relative to each other.

We were unable to resolve the cytoplasmic tail of F within the sub-tomogram average. To better visualize the position of F relative to the M lattice we modelled a straight rod through the center of the F trimers spanning across the membrane region. Figure 3F shows the rods (yellow) modelled into the sub-tomogram average along with models of RSV F timers fit into the map as viewed from the inside of the virion. This demonstrates that the two F trimers are centered over similar positions in the lattice between M-dimers. However, the center-to-center distance between the modelled F-trimers (Figure 3F) was measured to be 7.59 nm compared to the 8.33 nm distance between two M dimers (Figure 2G). This discrepancy could arise from the flexibility of the F-trimer stalk and cytoplasmic tails [49, 50]. The sub-tomogram average included density connecting the two trimers (Figure 3B,E). Though this was not sufficient to confirm a direct interaction, the presence of an interaction between the trimers would be consistent with the trimers having a closer center-to-center distance than the cytoplasmic tails.

### Sub-tomogram averaging higher-order organizations of F

As shown in Figure 1D, the organization of F on the virus was not limited to isolated pairs, but appeared to have higher order organization, such as the presence of rows (Figure 1D). Classification of the particles in our sub-tomogram average of the F pair revealed that neighboring trimers were frequently located in specific positions relative to the central pair. We then used these classes as initial references for sub-tomogram averaging to identify particles containing clusters of four trimers. Figure 4A,E,I,M show slices from four different sub-tomogram averages, each with two pairs of trimers. Each average included between two and six hundred sub-tomograms.

**Figure 4.**
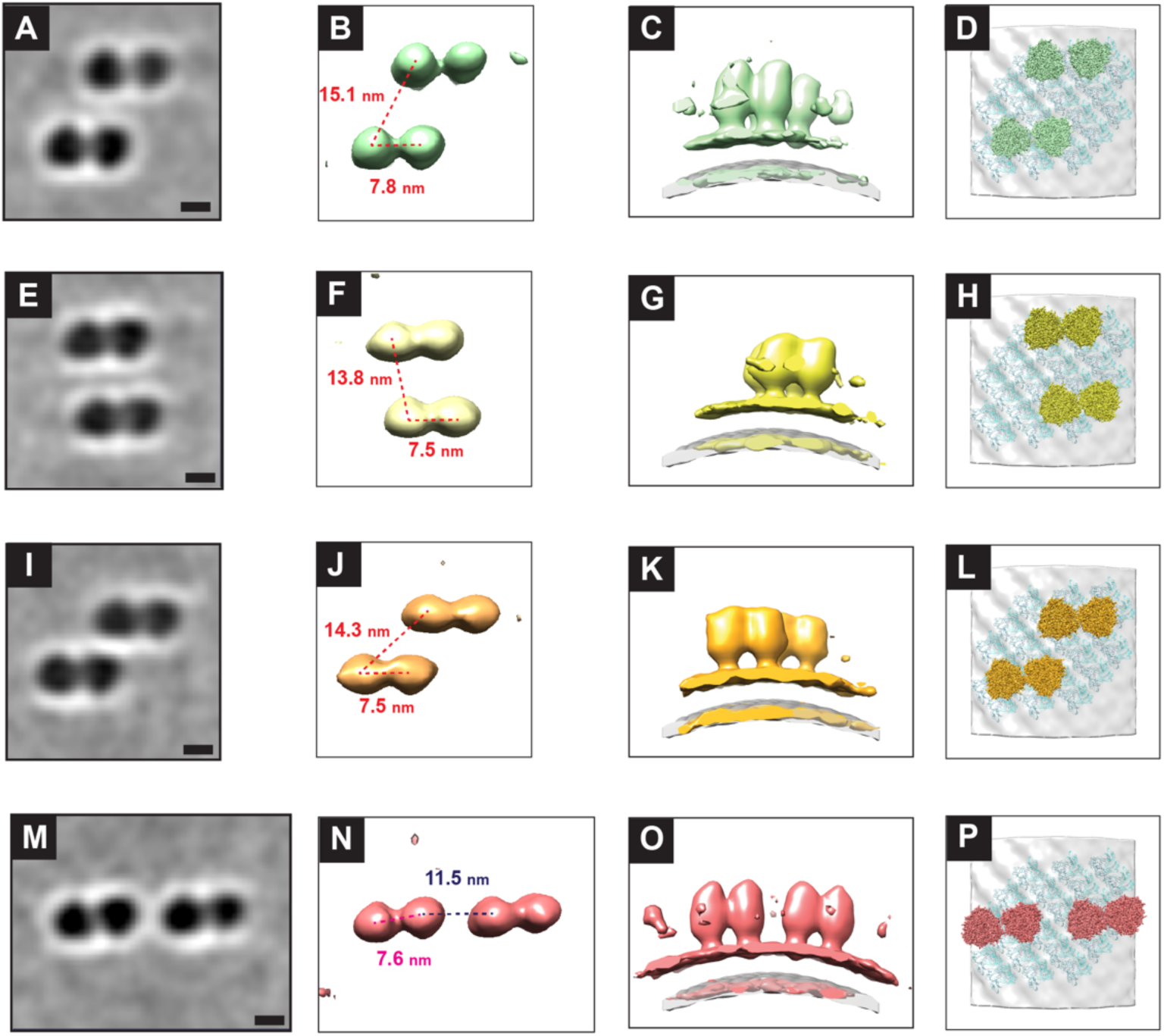
Sub-tomogram averages of RSV F pairs. A) Single slice from sub-tomogram average of four RSV F trimers (left). B) Isosurface of the sub-tomogram average in A (showing only density above membrane). Measurements between individual trimers are indicated with dotted lines. C) Isosurface of sub-tomogram average in A (green) aligned to the M lattice from the sub-tomogram average in Figure 2 (semi-opaque gray). D) Rotated view of C showing RSV F pre-fusion timer models (PDB: 5w23, green) fit into the sub-tomogram average of F in place of the isosurface. Models of the M-dimer (PDB: 4v23, cyan) were fit into the aligned density from the sub-tomogram average of the M lattice from Figure 2. E-P) As in (A-D) for different sub-tomogram averages with different distinct organizations of F trimers. 5 nm scale bar for (A),(E),(I),(M); volumes were lowpass filtered to 40 Å.

The anti-parallel and conserved orientation of the trimers can be seen in the sub-tomogram averages. To better visualize the trimer orientation and spacing, the pre-fusion F crystal structure was modelled into the sub-tomogram averages (Figure 4D,H,L,P). Using the center of the models, we were able to measure the distance and angles between trimers (Figure 4 B,F,J,N). The distance between individual trimers in the pair ranged from 7.5-7.8 nm, close to the 7.6 nm spacing measured in Figure 3. The slight variations in spacing between trimers and the rotation of individual trimers may be due to the limited resolution from the lower number of particles included in each average relative to the averages in Figures 2 and 3. We were able to fit the M lattice from the sub-tomogram average in Figure 2 to the averages in Figure 4 (Figure 4C,G,K,O). The allowed us to indirectly model the M dimer (PDB: 4v23) into the M lattice underneath the fit models for the F trimers (Figure 4D,H,L,P).

## Discussion

Matrix proteins have been identified as central organizers of virus assembly in negative-strand RNA viruses [20-23]. RSV and other *Pneumoviridae* family members are closely related to viruses of the *Paramyxovidae* family, including measles virus and Newcastle disease virus (NDV) for which the organization of matrix has been determined [29, 32]. For both viruses, sub-tomogram averaging was previously used to show that matrix is arranged in a lattice of dimers and the glycoproteins are similarly arranged in a lattice [29, 32]. However, given the smaller size of RSV M (28 kDa) and the fact that it was initially crystallized as a monomer [51], questions have remained about the organization of M in RSV. Our results from cryo-ET and sub-tomogram averaging of virions show that that M is arranged in a locally-ordered lattice of dimers with an apparent helical organization relative to the viral long axis. This is consistent with the reported organization in related viruses and prior studies that found M dimerization is required for the assembly of filamentous VLPs [25, 29, 32-34].

### Organization of RSV M

Model fitting of the RSV M crystal structure (PDB 4v23 [25]) into the sub-tomogram average reveals a tightly packed lattice of RSV M. The repeating M dimer lattice has an average spacing of 5.4 nm by 6.7 nm (Figure 2G). The orientation and spacing of this lattice is similar to the packing of the crystallized form of M (PDB:4v23) that had an asymmetrical unit consisting of an M dimer with unit cell dimensions of 52.3 Å, 79.2 Å, and 65.9 Å. The orientation of RSV M relative to the membrane in the sub-tomogram average matches what is expected from previous structural analysis of RSV M [51]. Conley et al. have recently reported a sub-tomogram average of the RSV M lattice with a similar orientation and organization to our results presented here [19].

Our results provide further evidence that lattice organization of matrix is broadly conserved across a number of negative-strand RNA virus families. In many cases the structure of matrix itself is conserved, crystal structures of RSV and NDV matrix proteins have an RMSD of only 3Å [32]. Lattice organizations of matrix proteins have been shown for a number of viruses with diverse virion morphologies including NDV [32], measles virus [52], rabies virus [53], vesicular stomatitis virus (VSV) [31], Ebola and Marburg viruses [33], influenza A virus [34] and others. VSV and rabies virus have bullet-shaped virions approximately 70 nm in diameter and 200 nm in length with relatively low variation (∼50 nm) in either dimension [31, 53]. Measles and NDV have ellipsoidal virions that vary in shape from spherical to near filamentous. The dimensions of measles virions can range from 50 nm to >500 nm [29, 30]. Ebola and Marburg have filamentous virions that vary in diameter by ∼20 nm, with lengths extending to many micrometers [33], while influenza A forms both spherical and filamentous virions [34, 54]. The actual organization of the lattices formed by viral matrix proteins is also quite variable. The matrix lattices of Measles and NDV have four-fold symmetry, VSV and rabies virus matrix exist as true helical assemblies, while other lattices formed by matrix retain two-fold symmetry with more limited helical character. The apparent helical pitch among those with two-fold and/or helical symmetry varies from close to zero for Marburg and rabies viruses [33, 53] to the 47° reported here for RSV.

RSV has filamentous virions that can vary in length by several micrometers and in diameter by more than two-fold (Supplemental Figure S4, [6, 14, 19]). Arrays of M can be seen lining the membrane of filamentous virions with a range of diameters, as well as along flatter segments of irregular viruses. However, regions of ordered M are often not present in areas of high membrane curvature such as the ends of filamentous particles, and at bends or branch points in irregularly-shaped virions [6, 7, 14]. Different curvatures of the lattice would require lattice flexibility supported by altered spacing between subunits. Flexibility in the M lattice could be achieved through plasticity of the dimer interface, which has been demonstrated through comparison of the two M dimer crystal structures [25]. There may be additional flexibility in dimer-dimer organization.

The presence of gaps or irregularities in the lattice could be an additional mechanism for achieving a range of local curvatures, but also suggests a limitation to the flexibility of the intact lattice. Similar irregularities have been observed in the matrix protein lattice of other viruses [32, 33, 55, 56]. We observed minor variations in measurements of the lattice spacing both between subunits in the sub-tomogram average and in the Fourier analysis from different virion segments. This variation may reflect flexibility of the M lattice, though limitations such as imperfect fitting of the model into the sub-tomogram average should also be considered. Using the 47º apparent helical pitch measured in our sub-tomogram average (Figure 2G) and subtracting 14 nm from the measured diameters (Supplemental Figure S4) to account for the spacing between the viral membrane and M lattice, the range for the radius of curvature at the level of the M lattice was calculated to be 91.3 nm to 203.1 nm (Supplemental Figure S4).

One potential mechanism for driving the diversity of virion morphology and lattice structure is the contribution of additional structural proteins. Ebola VP40 and measles virus M are sufficient to assemble filamentous VLPs [57, 58]. For Ebola, the expression of additional structural proteins alters both the VLP diameter and helical pitch of the VP40 lattice [33]. Nipah virus M, F, and to a lesser extent G are all independently capable of budding VLPs. Influenza matrix (M1) associates with, but is not required for VLP formation driven by Hemagglutinin and Neuraminidase. Purified RSV M can assemble into a uniform helical filament in the presence of specific lipids [59], but requires expression of P and the F cytoplasmic tail for assembly of filamentous particles in cells [36, 37]. Many viral matrix proteins have also been shown to interact with and/or coordinate the positioning of nucleoprotein [30-32, 34, 52, 53].

### Organization of RSV F

RSV F is a class I fusion protein that may also support cell attachment. Though direct interaction between RSV M and F has not been shown, evidence for an interaction between matrix and fusion proteins has been reported for RSV and related viruses [60-63]. The cytoplasmic tail of RSV F is essential for filamentous VLP assembly [61]. RSV F is present on virions in a prefusion state and undergoes an irreversible conformational change during membrane fusion into a post-fusion state. The transition to the post-fusion state can be induced by other factors, as well, such as heat [7]. Structures of the RSV F homotrimer have been solved in both the pre- and post-fusion states, though these structures lack the transmembrane segment and cytoplasmic tail [13, 50]. Sub-tomogram averaging of RSV F on virions has been used to distinguish between the pre- and post-fusion states of large numbers of individual trimers [8, 14].

Our sub-tomogram average in Figure 3 contains a pair of potentially interacting F trimers. Fitting of the prefusion-F trimer crystal structure confirmed that the paired densities on the virus surface visible in the tomograms are F trimers. The trimers are oriented relative to one another within a pair and the pairs share a common orientation relative to the virus. This was also seen in our sub-tomogram averages in Figure 4 with four F trimers each. While pairing of trimers could be explained by local interactions, the common orientation of pairs requires a model for long-distance regulation of F positioning on the virion. Additionally, the pairs of trimers are positioned in preferred positions relative to one another as shown in the sub-tomogram averages in Figure 4. All of our data support a model in which each F-trimer is positioned over an equivalent point relative to an underlying M-dimer. This may occur at an M dimer-dimer interface as indicated by the modelling in Figure 3F, however given the potential flexibility of the F stalk and cytoplasmic tail, further work is necessary to determine the precise position. An interaction between the F-cytoplasmic tail and the M lattice would be one possible mechanism. This model is further supported by the necessity of the F-cytoplasmic tail to assemble filamentous virus-like particles [61, 62].

Applying this model to interpret the distribution of F on the viral surface, F trimers can occupy positions in rows roughly perpendicular to the viral long axis with no unoccupied positions in between as in Figure 4M. The formation of pairs pulls the F trimer heads closer together causing them to appear unevenly spaced despite their regular spacing relative to the M lattice. Unoccupied positions in these rows further contribute to the appearance of uneven spacing. The sub-tomogram averages in Figure 4A and 4E represent pairs of F trimers with two unoccupied rows between them while the pairs in Figure 4I only have a single unoccupied row between them. This variation in the gaps between rows means that though the positions of the trimers are regulated by the underlying helical-like M lattice, it may not be possible to follow a single row of F-trimers over an entire helical turn around the virus.

The presence of F pairs raises many interesting questions as to whether this coordinated arrangement is functionally important for the virus. Close packing of F could affect antibody and/or receptor binding or it may promote fusion since the clustering of class I fusion proteins is known to occur during membrane fusion [64-66]. Pairing of F could also affect the stability of the F prefusion state [67, 68]. It is possible that the ordered rows of F pairs on the virions are one transient F arrangement that exists over a continuum of intermediate F trimer states. This overall organization of F trimers may evolve over time due to fluidity in the viral membrane and underlying M lattice, possible associations with secreted or membrane-associated G, or interaction with receptor or other cellular proteins. Furthermore, it is known that class I viral fusion proteins exist in a ‘metastable’ prefusion structural state, and that upon ‘triggering,’ which is caused by cascading interactions with another glycoprotein and/or cellular receptor and/or co-receptor, the protein undergoes an irreversible conformational change to the post-fusion state [13, 50, 68].

Until recently there has been limited structural information about the coordinated organization of fusion or attachment glycoproteins in either purified protein preparations or on intact virus particles. For example, negative stain TEM and cryo-ET of purified HPIV3 illustrated that HN, in the ‘heads down’ conformation, formed regular arrays on virions and that F was absent in these regions [69]. However, no lattice-like arrangement of pre- or post-fusion F alone or F colocalized with HN in the ‘heads up’ orientation was observed on the virions, indicating the transient nature of these glycoprotein arrangements. We have demonstrated that measles virus F is presented in an organized lattice with four-fold symmetry coordinated by an underlying lattice of M [29]. In addition, Xu et al. demonstrated by x-ray crystallography and EM that prefusion stabilized Nipah virus F trimers could form a semi-stable hexamer of trimers both *in vitro* and on virus-like particles (VLPs) [67]. Thus, validating the requirement for preserving and locking these classes of proteins into a single conformational state for intermediate to high-resolution structural analyses [50, 67, 70]. Though we did not observe hexagonal organization of RSV F on virions in this study, hexagonal lattice packing of RSV F on intact, filamentous virions has been reported for vaccine-candidate strains that were engineered with stabilizing mutations in F and reduced G levels [52]. This further supports the ability of RSV F prefusion trimers to assemble into varied states of higher-order oligomerization, potentially in a transient or condition dependent manner. Arrays of RSV F were also observed on irregular particles by Conley et al [19]. In kind, the fusion protein of influenza type C, HEF1, was shown to assemble as hexagonal arrays [71]. The inter-trimer interactions of F trimers within the Nipah F hexamers and HEF hexagonal array are similar to what we report for RSV F dimers (Figure 3B,E) providing evidence that the assembly of RSV F into pairs or hexamers may occur through interactions conserved between a number of viral class I fusion proteins.

Our sub-tomogram averaging from whole-cell cryo-ET of RSV virions shows that M is present as a tightly packed lattice of dimers with a helical-like arrangement in filamentous virions. Further, the RSV F glycoprotein is not randomly distributed on the virion, but frequently occurs as pairs of trimers and in specific positions consistent with coordination of F positioning by the underlying M lattice. Future studies will be needed to determine the nature of the structural contacts present between the M lattice and F as well as conditions under which RSV prefusion F trimers oligomerize and the functional role of these arrangements.

## Materials and Methods

### Cell culture and infection on TEM grids

BEAS-2B cells (ATCC CRL-9609) were cultured and maintained at 37°C with 5% CO_2_ in RPMI-1640 (Thermo Fisher Scientific) media supplemented with 10% fetal bovine serum (FBS, Hyclone) and 1x antibiotic antimycotic solution (100 units/mL penicillin, 0.1 mg/ml streptomycin and 0.25 μg/ml amphotericin B, Thermo Fisher Scientific). Vero cells (ATCC CCL-81) were maintained in DMEM (Thermo Fisher Scientific) supplemented with 10% FBS and 1x antibiotic antimycotic solution. Cells were released for passaging or seeding onto EM grids using 0.25% trypsin (Thermo Fisher Scientific). Cell counting was done using a hemacytometer following trypan blue staining.

### Respiratory syncytial virus and infection on TEM grids

The RSV strain rA2-mK+ strain was used for all experiments. RSV rA2-mK+ shares the same growth kinetics as its parent strain, A2, and expresses the fluorescent protein mKate2 during replication due to insertion of this gene at the 3’ end of the genome [72]. Viral propagation and fluorescent focal unit (FFU) titers were performed using Vero cells as described previously [6].

To provide extra support for cell growth 5-6 nm of carbon was evaporated onto Quantifoil Au 200 R2/1 or R2/2 grids (Quantifoil Micro Tools GmbH). Grids were then glow discharged and incubated in supplemented RPMI-1640 overnight in the cell incubator. Two to four grids were placed in a 30 mm glass bottom dish (MatTek Corp) and 7.5×10^4^ BEAS-2B cells were added in 2 ml of media. After overnight incubation (15-17 hours) the media was removed and the cells were infected with RSV rA2-mK+ at a multiplicity of infection (MOI) of 10. The grids were plunge frozen using a Cryoplunge 3 (Gatan, Inc.) 24 hours post infection. BSA gold tracer (10 nm, Aurion) was added immediately prior to freezing as a fiducial marker for tilt-series alignment.

### Cryo-electron microscopy and tomography

CryoEM images were collected on a Titan Krios 300 kV electron microscope (Thermo Fisher Scientific) equipped with a Gatan K3 direct-electron detector and BioQuantum energy filter (20 eV slit width, Gatan, Inc.). Tilt-series were collected bidirectionally from 0° in 2° increments from -60° to +60° with a total dose of ∼80e/Å^2^ using SerialEM [73]. The unbinned pixel size was 4.472 Å and the nominal defocus was -5 μm.

Frames were aligned with the built-in SerialEM algorithm or IMOD alignframes. Tilt-series were aligned and processed in IMOD [74] including CTF correction by phase flipping and erasing gold fiducials. Tomograms were reconstructed in IMOD using weighted back-projection. EMAN2 2.91 [75] was used for lowpass gaussian filtering of tomographic slices and calculation of the amplitude spectrum using the math.realtofft processor. The density profile plot was generated in Fiji 1.53 [76].

### Sub-tomogram averaging

All sub-tomogram averaging was done in PEET 1.15.0 [77, 78]. In order to generate an initial reference, thirty filamentous sections from 19 virions present in eleven tomograms were oriented as in Figure 1 with the long axis vertical and flat in the image plane. These sections were generated from tomograms binned by two. For sub-tomogram averaging of M: densities within the lattice were manually added as points and aligned and averaged with a limited search angle. Duplicate points and points with low correlation were discarded and this initial reference was then used for semi-automated particle picking. For semi-automated particle picking, the M layer of the virion segments was manually segmented and regularly spaced points were seeded throughout the segmented area. The seeded points were aligned to the initial reference allowing sufficient translation to overlap with the search range of neighboring points. Duplicate points and points with low correlation were discarded. The points and orientations were mapped back to the full unbinned tomograms and iterative rounds of reference refinement and alignment were performed to generate the final average shown in Figure 2. A mask with gaussian blur and missing wedge compensation were used during alignment. No lattice or other symmetry was applied. Particles were considered duplicates within 7 pixels (∼3 nm) thus particles from neighboring lattice points do overlap. The data was split into half-sets following alignment of the full set based on the initial thirty segments to ensure that any overlapping particles were in the same half-set. UCSF Chimera 1.15 [79] and Relion 3.1.3 [80] were used to generate a soft-edged mask around the M lattice. Relion was used to generate an FSC plot with the masked volume. The final number of particles in the M sub-tomogram average was 29,280. The reported resolution by fourier shell correlation (FSC) at the 0.5 criteria from the masked, corrected FSC curve was 20.78. The final sub-tomogram average was lowpass filtered to 21 Å in EMAN2.

Sub-tomogram averaging of the F pair started from the same thirty-oriented virion segments as M and followed a similar process. Points were picked manually to generate an initial reference using PEET, followed by seeding of regularly spaced points, semi-automated particle picking, and iterative rounds of reference refinement and alignment. UCSF Chimera and Relion were used to generate a soft-edged mask around the pair of F trimers. Relion was used to generate an FSC plot from the masked volume. The final number of particles in the F sub-tomogram average was 12,769. The reported resolution by FSC at the 0.5 criteria from the masked, corrected FSC curve was 23.85. The final sub-tomogram average was lowpass filtered to 24 Å in EMAN2.

For the sub-tomogram averages of high-order organizations of F we performed principle component analysis (PCA) and clustering in PEET to split the particles in the F-pair sub-tomogram average into multiple classes. The sub-tomogram averages from classes that represented unique organizations of F were used as an initial reference for multiple rounds of alignment, reference refinement, and classification on tomograms binned by 2 (8.944 Å/px). The averages in Figure 4A,E,I,M include 571, 318, 251, and 443 particles respectively. Isosurface rendering, model fitting, and measurements of sub-tomogram averages were performed using UCSF Chimera. Visualization of model points over the tomographic slice was done using the Place Object plugin [81].

## Data availability

All relevant data are available from the corresponding author upon reasonable request. All sub-volume averages have been deposited in the Electron Microscopy Data Bank (www.emdatabank.org) under the following accession numbers: XXXX

## Acknowledgements

This work was supported in part by the University of Wisconsin, Madison, the Department of Biochemistry at the University of Wisconsin, Madison, and public health service grants R01 GM114561 and U24 GM139168 to E.R.W. from the NIH. We are grateful for the use of facilities and instrumentation at the Cryo-EM Research Center in the Department of Biochemistry at the University of Wisconsin, Madison. The authors gratefully acknowledge use of facilities and instrumentation at the UW-Madison Wisconsin Centers for Nanoscale Technology (wcnt.wisc.edu), which is partially supported by the NSF through the University of Wisconsin Materials Research Science and Engineering Center (DMR-1720415). A portion of this research was supported by NIH grant U24 GM129547 and performed at the PNCC at OHSU and accessed through EMSL (grid.436923.9), a DOE Office of Science User Facility sponsored by the Office of Biological and Environmental Research. Molecular graphics and analyses performed in part with UCSF Chimera, developed by the Resource for Biocomputing, Visualization, and Informatics at the University of California, San Francisco, with support from NIH P41-GM103311.

## Figures and Figure Legends

**Supplemental Figure S1.**
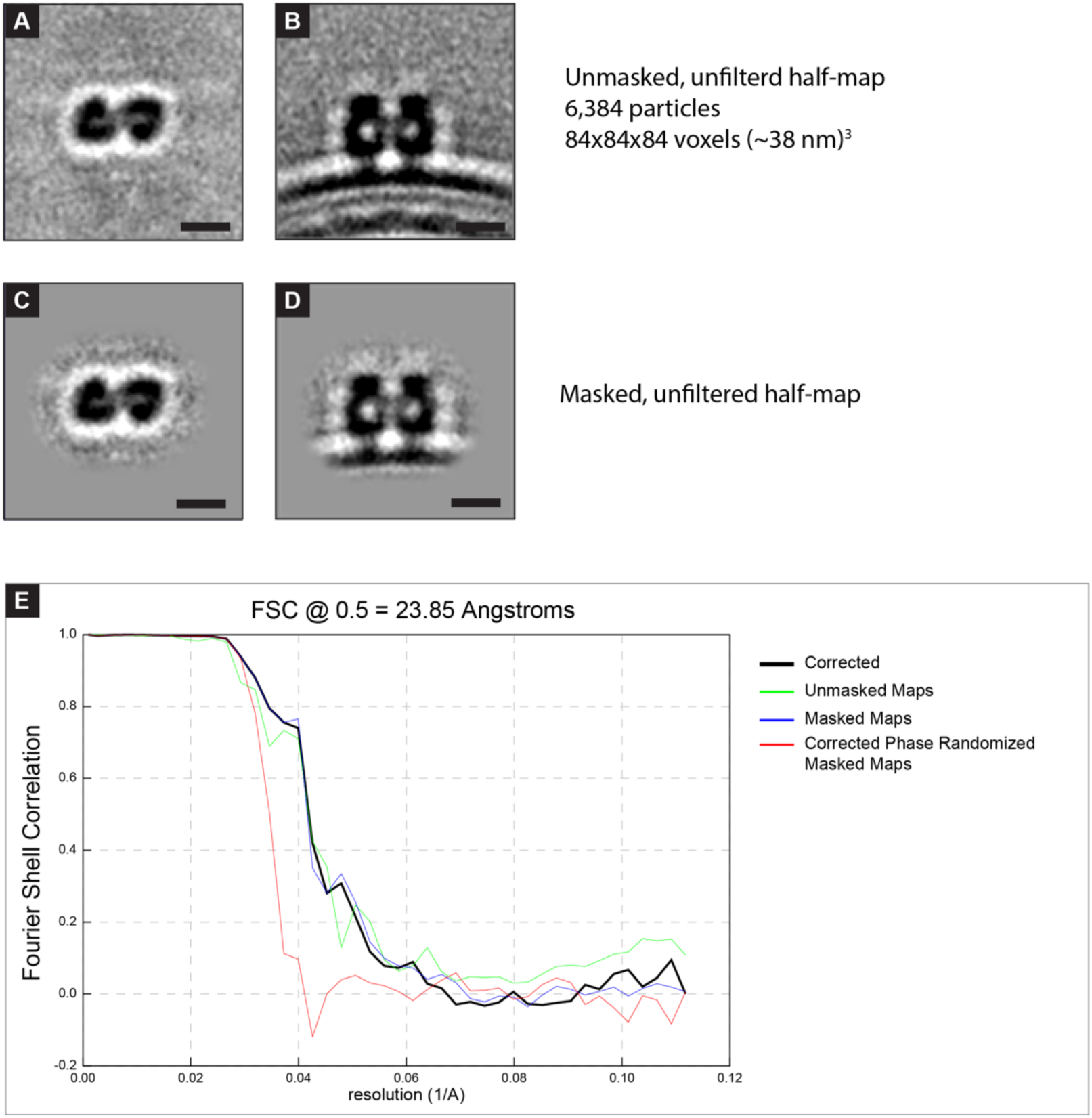
Analysis of the sub-tomogram average of the RSV F pair. A,B) Slices from sub-tomogram average containing half of the particles used in the final average of the RSV F pair. C,D) As in (A) with a soft-edged mask applied around the pair of F-trimers. This masked half-map and the corresponding half-map (not shown, remaining 6,385 particles) were used for the FSC calculation in E. E) Fourier shell correlation plot generated from the two half-maps. Resolution is reported using FSC 0.5 of the masked corrected plot (black line) because all particles were mutually aligned prior to generation of half-maps. 5 nm scale bar in (A),(B),(C),(D).

**Supplemental Figure S2.**
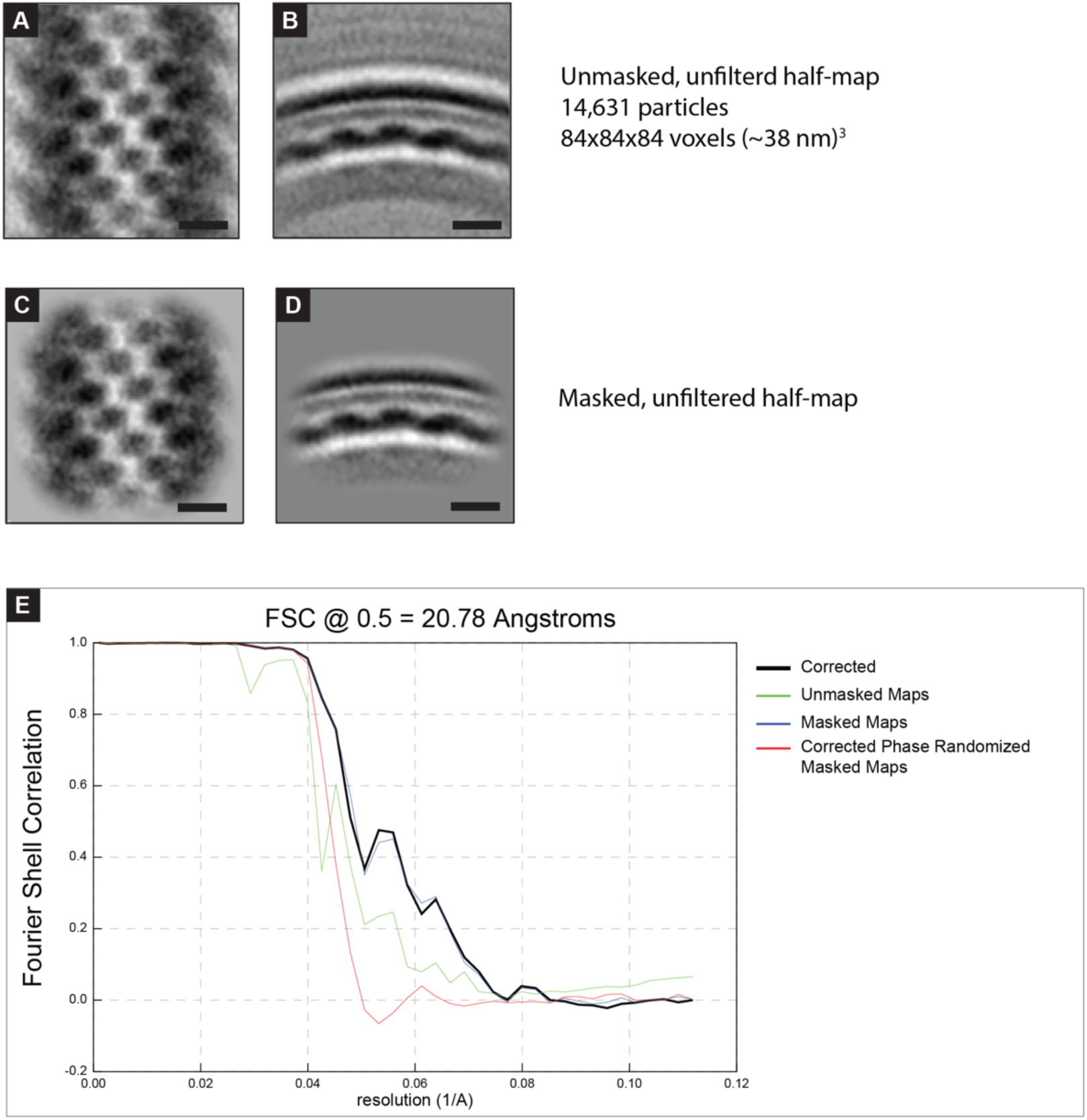
Analysis of the sub-tomogram average of RSV M lattice. A,B) Slices from sub-tomogram average containing half of the particles used in the final average of the RSV F pair. C,D) As in (A) with a soft-edged mask applied around the M lattice. This masked half-map and the corresponding half-map (not shown, remaining 14,649 particles) were used for the FSC calculation in E. E) Fourier shell correlation plot generated from the two half-maps. Resolution is reported using FSC 0.5 of the masked corrected plot (black line) because all particles were mutually aligned prior to generation of half-maps. 5 nm scale bar in (A),(B),(C),(D).

**Supplemental Figure S3.**
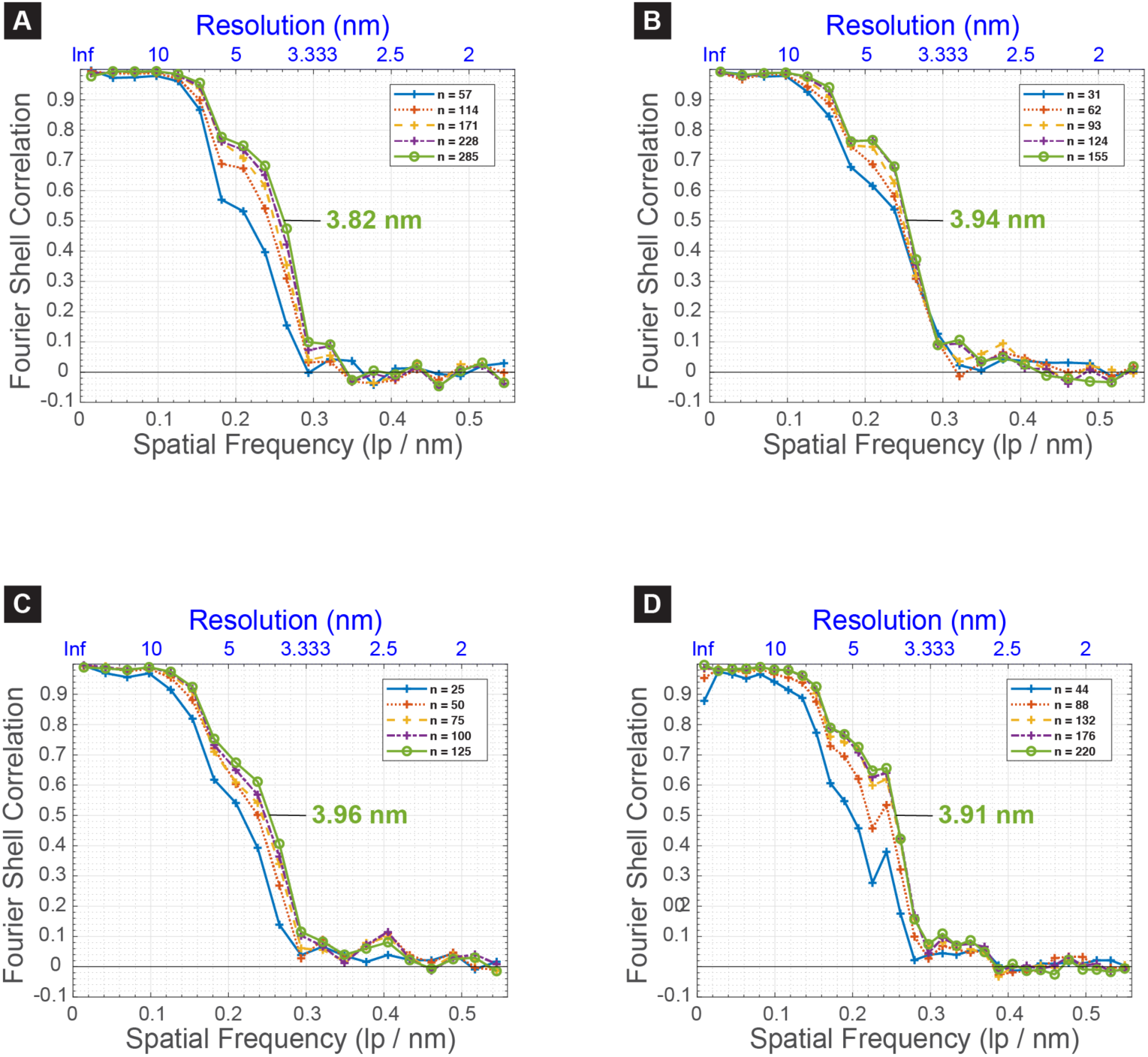
FSC curves for sub-tomogram averages of higher-order F organizations. A) FSC curves for the sub-tomogram average in Figure 4A from all particles (solid green line) or subsets of the data as calculated by PEET calcFSC from mutually aligned particles. The number of particles included in each half-set is indicated in the inset legend. The FSC at 0.5 for all particles is also indicated on the chart. B-D) As in A, for the sub-tomogram averages in Figure 4E,I, and M respectively.

**Supplemental Figure S4.**
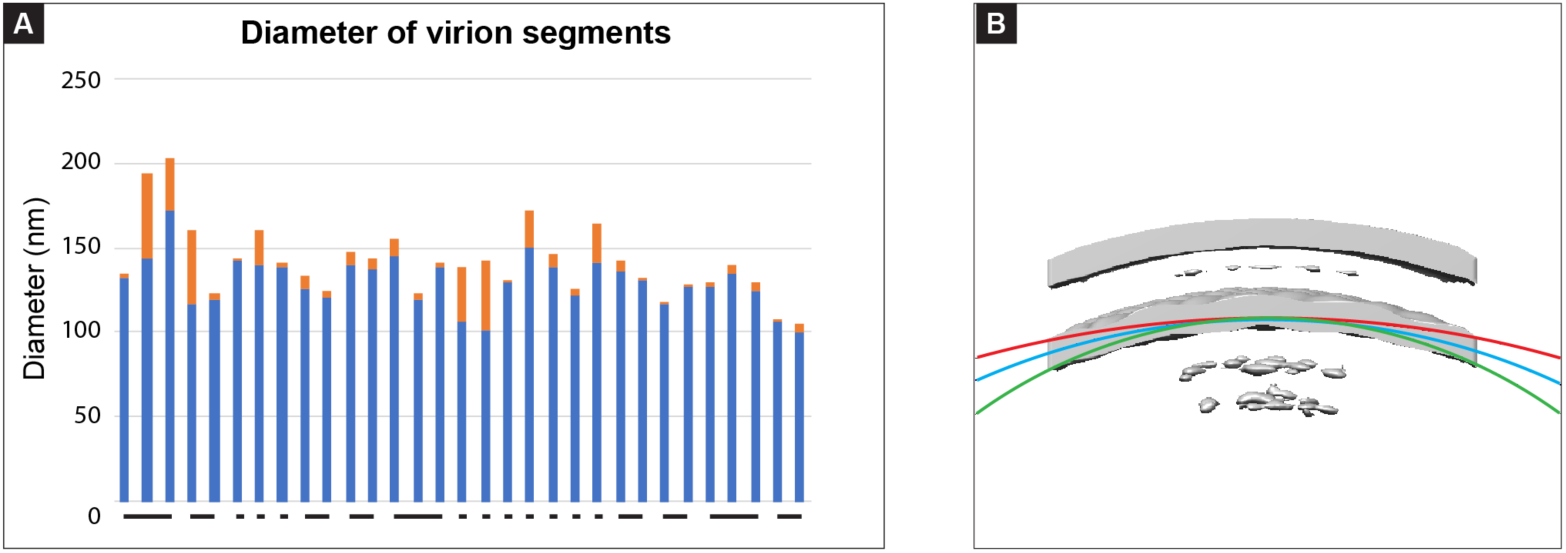
Diameter of RSV virion segments used for sub-tomogram averaging. A) Measurements of the virion diameter (membrane-to-membrane) near each end of the virion segment are graphed with the blue and orange bars representing the smaller and larger diameter of the segment respectively. Individual segments from the same virion are indicated by a connecting line beneath the bars in the graph. B) Arcs with curvature equivalent to circles with the measured minimum, maximum, and average virion diameter less 14 nm to account for the spacing between the membrane and M lattice are overlaid onto the M lattice sub-tomogram average. Red - 189.6 nm; Blue - 121.6 nm; Green - 85 nm.

## Notes

### Competing Interest Statement

The authors have declared no competing interest.

